# Single-color Fluorescence Lifetime Cross-Correlation Spectroscopy *in vivo*

**DOI:** 10.1101/2020.01.23.917435

**Authors:** M. Štefl, K. Herbst, M. Rübsam, A. Benda, M. Knop

## Abstract

The ability to quantify protein concentrations and to measure protein interactions *in vivo* is key information needed for the understanding of complex processes inside cells, but the acquisition of such information from living cells is still demanding. Fluorescence based methods like two-color fluorescence cross-correlation spectroscopy can provide this information but measurement precision is hampered by various sources of errors caused by instrumental or optical limitations such as imperfect overlap of detection volumes or detector cross-talk. Furthermore, the nature and properties of used fluorescent proteins or fluorescent dyes, such as labeling efficiency, fluorescent protein maturation, photo-stability, bleaching and fluorescence brightness can have an impact.

Here we take advantage of lifetime differences as a mean to discriminate fluorescent proteins with similar spectral properties and to use them for single-color fluorescence lifetime cross-correlation spectroscopy (sc-FLCCS). By using only one excitation and one detection wavelength, this setup avoids all sources of errors resulting from chromatic aberrations and detector cross-talk. To establish sc-FLCCS we first engineered and tested multiple GFP-like fluorescent proteins for their suitability. This identified a novel GFP variant termed slmGFP (short lifetime monomeric GFP) with the so-far shortest lifetime. Monte-Carlo simulations were employed to explore the suitability of different combinations of GFP variants. Two GFPs, Envy and slmGFP were predicted to constitute the best performing couple for sc-FLCCS measurements. We demonstrated application of this GFP pair for measuring protein interactions between the proteasome and interacting proteins and for measuring protein interactions between three partners when combined with a red florescent protein. Together, our findings establish sc-FLCCS as a valid alternative for conventional dual-color(dc)-FCCS measurements.

**STATEMENT OF SIGNIFICANCE:** The quantification of protein concentrations and protein-protein interactions *in vivo* is a crucial information needed for the understanding of complex processes inside cells. Determination of such information is unfortunately still challenging. Fluorescence-based method like fluorescence cross-correlation spectroscopy (FCCS) is the only method which provides this information *in vivo* and almost in the real time, however it suffers from limitations caused by experimental setup and biological origin of fluorescent proteins. We present single-color fluorescence lifetime cross-correlation spectroscopy as an alternative to FCCS, which uses the information of fluorescence lifetime to overcome some of these limitations. We challenged the method and determined its advantages and limitations and demonstrated the applicability of the method on the proteins of yeast proteasome.

## INTRODUCTION

The proteome of a cell is a complex mixture of millions of protein molecules of thousands of different species, all engaged in various types of interactions, from very transient ones to stable protein complexes. The fraction of an individual protein that is engaged in a functional interaction depends not only on the parameter that regulate the interaction, but also its own concentration and the concentration of its interaction partner(s) and of competing binding factors. Therefore, precise determination of protein *in vivo* concentrations, together with reliable measurements of association and dissociation constants, provides the necessary information to understand the dynamics of such a system. Methods such as immunoprecipitation or *ex vivo* studies using purified components(1, 2) provide qualitative information about the biochemical properties of individual proteins and their interactions. However, they do not necessarily explain the behavior of proteins in the crowded cellular environment with its many constituents. Here, interactions with small molecules and regulatory activities can exert major influences on protein-protein interactions. To overcome this, a number of methods have been developed to study protein-protein interactions in the context of the complex environment of the cell, either using crude protein extracts from lysed cells (Biacore(3)), or using indirect *in vivo* strategies employing functionalized reporter molecules (such as the ‘Two-hybrid’ and ‘Anchor away’ techniques(4, 5)). For a direct *in vivo* assessment of protein-protein interactions fluorescence microscopy can be used to monitor the proximity of molecules using FRET(6, 7) or super-resolution methods(8–10). For mobile and dynamic proteins, it is furthermore possible to estimate their interactions by quantification of co-mobility. One representative of these latter methods is dual color fluorescence cross-correlation spectroscopy (dc-FCCS)(11, 12). This method employs fluorescently labelled species of the molecules under investigation such as proteins tagged with different fluorescent protein reporters(13–15) or organic fluorescent dyes(16–19), and it analyzes the fluorescence fluctuations that result from the movement of the labelled molecules in- and out-of a specified confocal detection volume. Statistical analysis of the fluctuations that result from one or several fluorescently labelled species is termed fluorescence correlation analysis and provides information about the concentration and diffusive behavior (i.e. the diffusion coefficient) of soluble proteins. When conducted for two protein species simultaneously, each labelled with a different fluorophore, dc-FCCS enables the quantification of the fraction of both species that exhibit co-diffusion(20). In contrast to FRET, dc-FCCS provides reliable conclusions about the existence or the absence of a protein-protein interaction, since the obtained information is independent on the steric arrangement of the fluorophore. A major drawbacks of fluorescence fluctuation measurements however are the many sources of uncertainty associated with the analysis of the data. These originate from constraints imposed by the small measurement volumes of diffraction limited high NA optical systems, the heterogenous optical properties of the cellular interior where differences in the refractive indices of different cellular structures influence the shape of the detection volume, and a limited number of labelled species present in living cells. The situation is even further complicated by the *in vivo* properties of fluorescent proteins, e.g. slow maturation of their fluorophore or protein folding. Together with bleaching, this causes that not the entire population of the protein of interest is fluorescent. In addition, photophysical properties of the fluorophores, such as blinking or low quantum yield need to be considered(21).

Measurements via dc-FCCS make use of different fluorophores with their specific spectral characteristics. These measurements require different wavelengths for emission and detection of the signals in order to discriminate the fluorophores from each other. Such dual color measurements are associated with additional errors that are caused by the different sizes of the detection volumes in each channel, and by light scattering inside the cells that might affect the shape and size of the detection volume in a wavelength specific manner. This often yields an error that is influenced by the situation present in an individual cell and that is very difficult to correct for. In addition, dual color measurements suffer from bleed-through of the emitted photons from one fluorophore into the detection channel of the other fluorophore. This bleed-through results in an aberrant cross-correlation signal where the error associated with bleed-through correction directly limits the sensitivity by which weak protein-protein interactions can be quantified. In a typical situation using endogenously expressed proteins this limits the dynamic range for K_D_ measurements to values below 500 – 1000 nM(22).

Therefore, it is difficult to obtain reliable *in vivo* estimates of protein concentration and protein-protein interactions from a dc-FCCS measurement conducted in an individual cell. To address these limitations, many measurements performed in a representative population of cells in combination with statistical analysis of the data are required to obtain reliable and reproducible estimates of the desired parameter(22).

To improve the reliability of individual FCCS measurements it is desired to reduce the number of correction factors that are needed to analyze the fluorescence fluctuation data. To eliminate the volume overlap problem, it is possible to use fluorescent proteins with the same excitation wavelength, but different Stokes shifts resulting in well separated emission spectra. We developed the optimized long Stokes shift fluorescent protein mKeima8.5 and demonstrated that it can be used for FCCS measurements using a single excitation wavelength(23).

Furthermore, cross-correlation artifacts caused by bleed-through of the emission of one fluorophore into the detection channel of the other can be avoided by pulsed interleaved excitation (PIE) approach, where the green and red fluorophores are sequentially excited by the different lasers and the photons are distinguished based on their arrival time with respect to the laser pulse(24).

Here we now explore whether differences in the fluorescence lifetimes of different green fluorescent proteins (GFPs) can be used to eliminate the volume overlap and bleed-through problem in *in vivo* FCCS at the same time. For our analysis we employ the principle of fluorescence lifetime cross-correlation spectroscopy (FLCCS). FLCCS is a modification of FCCS, whereby a pulsed laser is used for fluorophore excitation and where each detected photon is weighted by a fluorescence lifetime, fluorophore-specific component based on its arrival time in relation to the excitation pulse. This enables to ‘filter’ the photons in each detection channel(25–27) and to statistically eliminate photons that are not emitted from the investigated fluorophore. In principal, with FLCCS it should also be possible to discriminate fluorophores with very similar spectral properties, provided that the lifetime histograms of their emitted photons are significantly different. With this a single color FLCCS (sc-FLCCS) setup could be used for fluorescence cross-correlation experiments, thereby eliminating the requirement for correction of two major sources of errors simultaneously: bleed-through and volume overlap.

Towards establishing sc-FLCCS, we evaluated first a broad range of GFP variants in order to identify suitable candidates with very short or very long fluorescence lifetimes. We use simulations, proof of principle *in vivo* measurements with synthetic constructs and real *in vivo* measurements with endogenously tagged proteins to challenge the method and to probe its limits. Our results indicate that sc-FLCCS is feasible and that it can be used to assess the interactions of multiple proteins *in vivo*.

## MATERIALS & METHODS

### Yeast strain construction and cell growth

All yeast strains were constructed using standard procedures as previously described(28) and validated by fluorescence intensity measurements, colony PCR and in some cases by sequencing. Prior to FCS measurement, yeast cultures were grown in synthetic complete medium (2% glucose) at 30°C (230 rpm) over night until saturation, diluted to OD_600_ ~ 0.1 and grown again to OD_600_ ~ 0.5 (30°C, 230 rpm). Then, the cells were immobilized on the glass surface of the microscopy plates (Greiner Sensoplate™, Greiner Bio-One GmbH, Austria) using Bio-conext (PSX1055, UCT Inc., USA) and Concanavalin A(22) and covered by Low Fluorescence medium (SC medium w/o riboflavin and folic acid). FCCS/FLCCS measurement was performed within the next 2-3 hours by pointing the laser to the cytoplasm of budding cells.

### Immunoblotting (Western blotting)

Whole-cell extracts were prepared using the sodium hydroxide/trichloracetic acid method(29). Proteins were further separated by SDS-PAGE using a 12 % polyacrylamide separating gel and transferred onto nitrocellulose membrane (XCell II Blot Module, Invitrogen). The membrane was incubated overnight with rabbit polyclonal primary anti-GFP antibodies (Abcam, ab6556). Peroxidase-conjugated goat anti-rabbit antibodies (111-035-003, Dianova, Germany) were used as secondary antibodies for detection. The visualization was performed on LAS-4000 imaging system (GE Healthcare Life Sciences).

### Monte-Carlo Simulations

As for Monte-Carlo simulations we extended the work of Wohland et al.(30) by adding excited state lifetime information and multiple diffusing components. The generated data were saved in TTTR data format, particularly in .pt3 files. Random positions of two types of particles (components), which differed in the excited state pattern and/or concentrations, were initially generated and then changed in each simulation step according to 2-dimensional Brownian diffusion in a 6 μm square simulation box with periodic boundaries. The diffusion coefficient was fixed to the value of 2.25 μm^2^s^-1^. The detection volume was approximated by a Gaussian profile with a beam waste radius of 300 nm. The simulations were run with 100 ns time steps, corresponding to 10 MHz repetition rate, with a sampling time of 32 ps per TCSPC channel, each time trace was 180 s long. A TCSPC channel for every generated photon was chosen by random selection from a look-up table corresponding to component’s excited state pattern (exponential with different lifetimes, Gaussian with different peak position or experimental). The simulated molecular brightness was 100 kHz per molecule, data with lower molecular brightness were generated from this 100 kHz data set by proportional random deletion of photons for the given component to save simulation time.

### Fluorescence Cross-Correlation Spectroscopy

Fluorescence correlation spectroscopy(11, 12) is based on the statistical analysis of the time-scale intensity fluctuations *I(t)*. Such dependence is described by the normalized auto- and cross-correlation functions *G_AC_* and *G_CC_* which are defined as

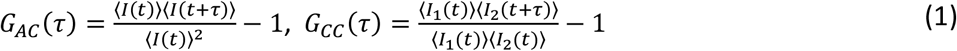

*τ* is the time lag, angle brackets denote the averaging over all possible values of time *t* and *I_1_(t)* and *I_2_(t)* are the intensity fluctuations in two different detection channels. In the case of Brownian motion in a three-dimensional Gaussian detection volume, assuming forbidden intersystem crossings and two species of molecules with distinct diffusion characteristics the auto-correlation function can be described by the following equation:

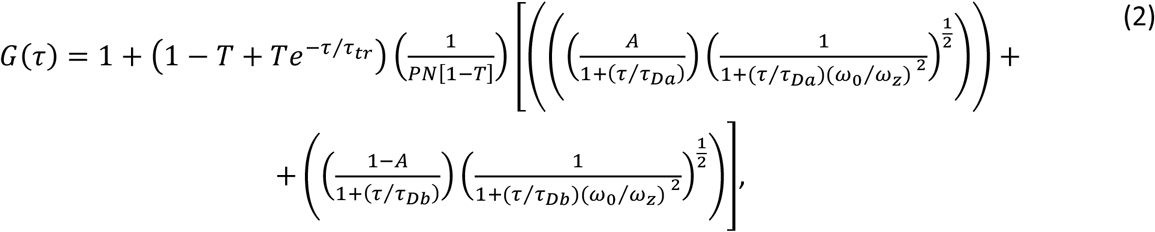

where *T* and *τ*_0_ are the contribution and kinetics of intersystem crossing, *PN* corresponds to the number of particles in the detection volume. *τ*_Da_ and *τ*_Db_ correspond to the average times of diffusing species <u>a</u> and <u>b</u>, for which the fluorescence molecules stay in the detection volume. *A* corresponds to the relative amplitude of the autocorrelation function with diffusion time *τ*_Da_ and is given by 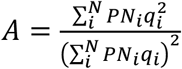, where *q_i_* corresponds to the molecular brightness of the *i-th* specie. The *ω*_0_ and *ω*_z_ are the spatial parameters of the detection volume. Thus, the profile of the auto-correlation function bears the information about the concentration and diffusion properties. The concentration can be calculated as:

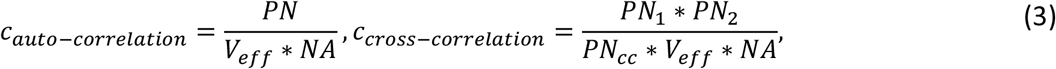

where *PN_1_, PN_2_* and *PN_cc_* correspond to the number of particles of individual species and their complex, *V_eff_* is the effective confocal volume and *NA* is the Avogadro number. The interaction between two proteins of interest can be represented by the dissociation constant which is defined as:

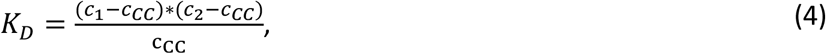

where *c_1_, c_2_* and *c_CC_* are the concentrations of species in channel *1, 2* and their complex.

### Fluorescence Lifetime Correlation Spectroscopy

Fluorescence lifetime histograms are measured by time correlated single photon counting, where the time axis is divided into small parts (bins). The width of the bin depends on the time resolution. The mathematical expression for the overall fluorescence histogram *I_j_* is:

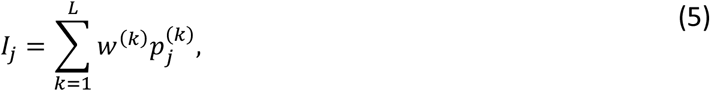

where *j* corresponds to the bin number, *L* to the number of decay components indexed by *k*. *w^(k)^* is the amplitude of the photon count contribution of the *k*th species and 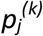 is the fluorescence decay pattern of *k*th specie alone, which is equal to its fluorescence lifetime histogram and is usually measured separately. In FLCCS(25–27, 31) the normalized auto-correlation function is described by the following expression:

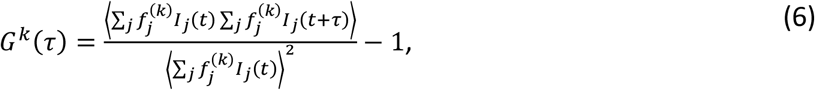

meaning that each photon is multiplied by a statistical filter *f_j_*, which corresponds to the fluorescence decay component *k*. The formula for the weighting factor 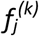 is:

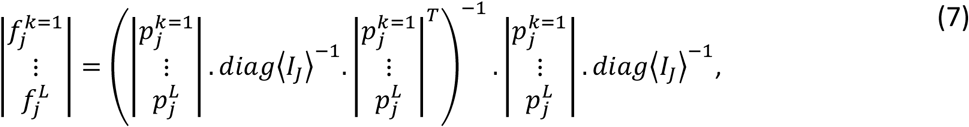

where the dot ·, superscript *T* and *^-1^* denote matrix multiplication, transposition and inversion.

### Microscope setup

FCCS/FLCCS were measured using a Nikon TiE body combining an inverted wide field fluorescence microscope with a confocal MicroTime 200 unit (PicoQuant, Germany) for time-resolved photon counting. Picosecond pulsed laser diode head (LDH-D-C-485; Picoquant, Germany) with a 20MHz repetition rate or a single frequency CW diode pumped laser (Cobolt Jive™ 561nm, Cobolt, Sweden) were used for the excitation of green and red fluorophores. The collimated laser beam was coupled into an optical fiber for optical cleaning and then reflected using a beam splitter (zt 488/561rpc; AHF Analysentechnik AG, Germany) into the inverted microscope body (Nikon Eclipse Ti; Nikon Instruments Europe B.V., Netherlands). The sample was illuminated using a water immersion objective (Nikon, Plan Apo IR, 60x/NA 1.27; Nikon Instruments Europe B.V., Netherlands) and the same objective was used for collection of the fluorescence light. Emitted light passed through the 50 μm pinhole, band-pass emission filters (ET525/50m and ET632/60m; Chroma Technology Corporation, VT, USA) and was detected by a τ-SPAD single photon avalanche photodiode (PicoQuant, Germany). Low laser intensities (< 5 μW for 485 nm and 561 nm) were used to prevent photobleaching and pile-up effect (in case of 485 nm excitation). All data were measured at 19 °C. The size of the detection volume was determined using calibration dyes Atto 488 (D = 390 μm^2^s^-1^, T = 19 °C) and Atto 565 (D = 390 μm^2^s^-1^, T = 19°C). Volume overlap was determined using double labelled *in vitro* FCCS standards for 488-543 nm (iba, Göttingen, Germany).

### Analysis pipeline

All experimental data was analyzed using a custom-made data analysis pipeline (Fig. S1 in the Supporting Material) written in Matlab (MathWorks, Natick, MA, USA). For single wavelength analysis using two different green FPs with different lifetime histograms, abbreviated as **sc-FLCCS**, we first measured fluorescence lifetime histograms of pure individual FPs (using protein fusions to the endogenous gene) and use them as reference patterns. Second, the reference patterns were loaded together with the filtered lifetime histogram and corresponding weighting filters were calculated according to the Eq. 7 (Fig. S2a). In the case of double color excitation (excited by pulsed 485nm laser and continuous 561nm laser), abbreviated as **dc-FLCCS** for dual color FLCCS, lifetime filtering was used to correct for the bleed-through of the photons which were excited by the pulsed 485nm laser and detected in the red channel. The weighting filters were calculated from the fluorescence lifetime histograms (Eq. 7) detected in the green channel. The ‘positive’ filter was used for filtering from the green detector and complementary ‘negative’ filter was applied to the red detector (Fig.S2b). Further mathematical operations (e.g. corrections and correlations) were common to both, the sc-FLCCS and the dc-FLCCS.

Next, we corrected for photo-bleaching by dividing the overall intensity time trace into a set of shorter time intervals (3 s intervals in our case), which were correlated individually (Eq. 6)(32). The final correlation curve was derived by averaging the multiple short-interval based correlation curves. Resulting auto- and cross-correlation curves were fitted with the model described in Eq. 2. Finally, the concentrations and dissociation constants were calculated (Eq. 3 and Eq. 4).

## RESULTS

### Characterization of green fluorescence proteins

To establish sc-FLCCS we first aimed to identify fluorescent proteins with optimal properties. To this end there are many publications that report different GFPs, however only for few of them information about the fluorescence lifetime is available. To obtain this information we selected 15 different GFP variants (Table 1): Envy and Ivy, NowGFP, GFPgamma, myeGFP (monomeric yeast enhanced GFP), sfGFP (superfolder GFP with and without the F64L mutation for improved folding at 37°C) and circular permutations of sfGFP (cp3, cp7 and cp8), no-sfGFP (sfGFP without the superfolder mutation), mNeonGreen, Clover (with and without the F64L mutation for improved folding at 37°C) and slmGFP (a new GFP variant combining different mutations (33–35)). We furthermore used yeast codon optimized variants of the corresponding genes for their expression in yeast to test their fluorescence lifetimes and other properties (Table 1). This identified lifetimes in the range of 1.8 ns to 3.9 ns, with slmGFP exhibiting the shortest lifetime of all. Using cells without fluorescent protein expression we also observed the weak cellular autofluorescence with a highly multi-exponential lifetime in the range of 2.4 ns. This indicates multiple sources for this autofluorescence, in particular riboflavins, a common source of autofluorescence in yeast(36). For further analysis we selected the three GFPs with the shortest and the four GFPs with the longest lifetimes for further evaluation (Fig. 1a, bold rows in Table 1).

**Fig. 1.**
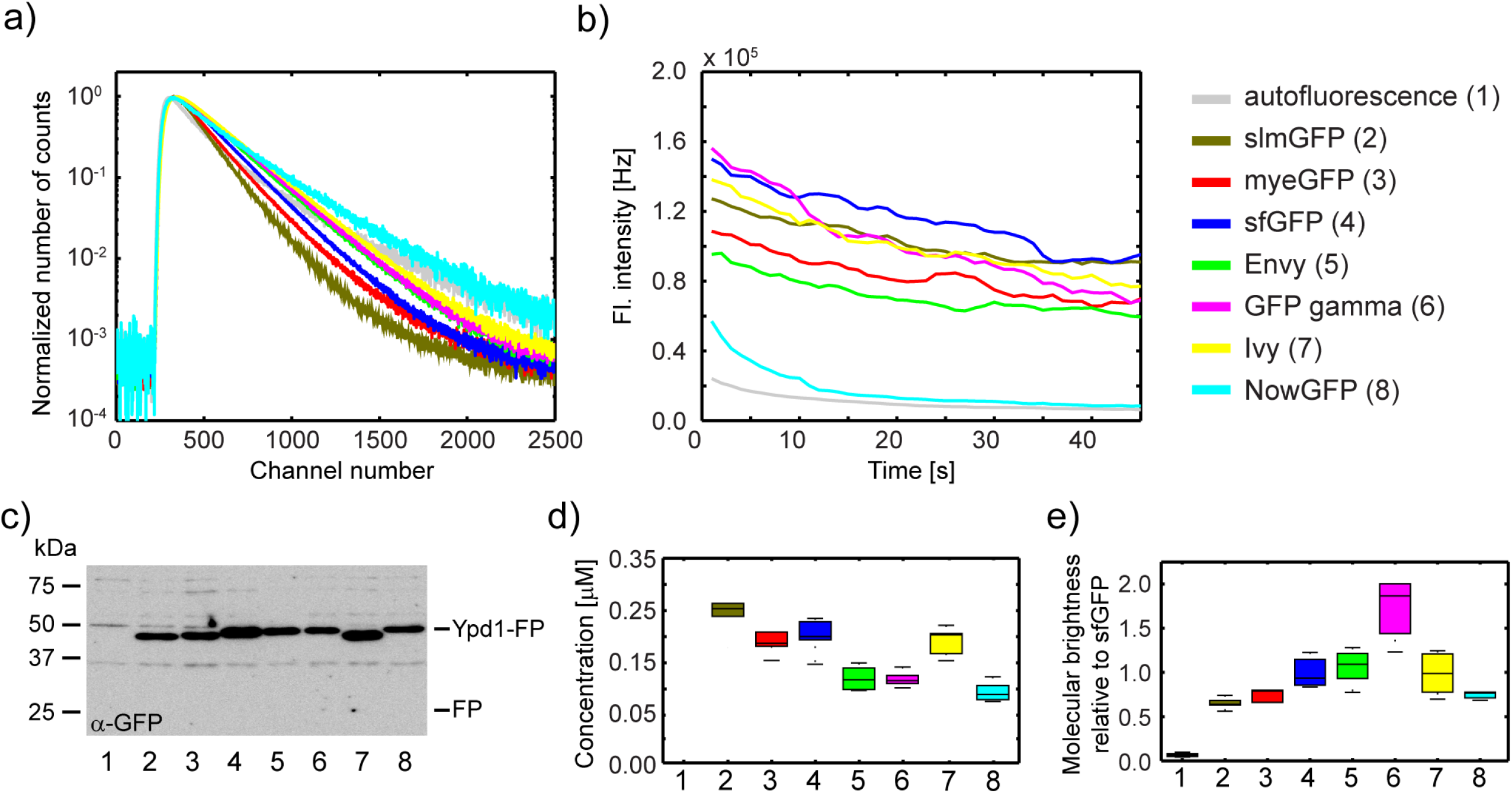
Fluorescence properties of selected green fluorescence proteins. Different fluorescence proteins (color coding/numbering is consistent throughout the whole figure) were expressed in yeast as an endogenously expressed C-terminally tagged Ypd1 fusion protein. **(a)** Fluorescence lifetime histograms normalized to the maximum of each curve. **(b)** Photobleaching curves. Raw fluorescence intensity traces which were measured for 45 s. Presented data in a and b correspond to average traces from 5 measurements. **(c)** Immunoblot and detection of the Ypd1-GFP fusion proteins using anti-GFP antibodies. Please note that this does not allow direct comparison of protein levels, since different GFP variants may cover different range of epitopes recognized by the polyclonal anti-GFP antibodies. No degradation products were detected, indicating that no free GFP is present inside the cells(40). **(d)** Absolute protein concentrations as determined by FCS. **(e)** Fluorescence molecular brightness normalized to the mean of Ypd1-sfGFP. Data in (d) and (e) are the result of 5 measurements each.

**Table 1.** Summary of fluorescence properties of various green fluorescence proteins. Proteins which were selected for further studies are indicated in bold. Fluorescence stability is defined as the percentage of fluorescence remaining after 20 s of illumination (corrected for the autofluorescence). Molecular brightness (as defined in the text) and fluorescence stability values correspond to the mean of 5 measurements per strain. Errors are expressed by standard deviations. Used abbreviations: not determined (n.d.), Absorption (Abs), Emission (Em), arbitrary units (a.u.).

| Protein | Fluorescence lifetime [ns] | Abs<sub>max</sub> [nm] | Em<sub>max</sub> [nm] | Molecular brightness [a.u.] | Fluorescence stability [a.u.] | Maturation [min] |
| --- | --- | --- | --- | --- | --- | --- |
| Autofluorescence | 2.40 ± 0.20 |  |  |  |  |  |
| **slmGFP** | **1.80 ± 0.15** | **501** | **513** | 0.65 ± 0.07 | **0.97 ± 0.13** | **4\*(37)** |
| **myeGFP Ref(22)** | **2.00 ± 0.18** | **501** | **513** | 0.76 ± 0.41 | **0.94 ± 0.19** | **n.d.** |
| **sfGFP Ref(38)** | **2.50 ± 0.14** | **487** | **511** | 1.00 ± 0.18 | **0.90 ± 0.07** | **6(39)** |
| sfGFP_cp8 Ref(40) | 2.52 ± 0.20 | 487 | 510 | n.d. | 0.77 ± 0.14 | n.d. |
| sfGFP_cp7 Ref(40) | 2.72 ± 0.11 | 486 | 510 | n.d. | 0.87 ± 0.06 | n.d. |
| sfGFP_cp3 Ref(40) | 2.78 ± 0.04 | 487 | 510 | n.d. | 0.84 ± 0.10 | n.d. |
| no-sfsfGFP Ref(40) | 2.84 ± 0.06 | 489 | 511 | n.d. | 0.90 ± 0.08 | n.d. |
| mNeonGreen Ref(41) | 2.90 ± 0.12 | 506 | 517 | 1.71(41) | 0.80 ± 0.27 | <10(41) |
| Clover (F64L) Ref(40) | 3.00 ± 0.03 | 506 | 517 | n.d. | 0.67 ± 0.12 | n.d. |
| Clover Ref(42) | 3.00 ± 0.09 | 505 | 517 | 1.56(41) | 0.69 ± 0.37 | 30(40) |
| sfGFP (L64F) Ref(40) | 3.10 ± 0.01 | 487 | 511 |  | 0.78 ± 0.09 | n.d. |
| **Envy Ref(43)** | **3.23 ± 0.02** | **486** | **509** | 1.06 ± 0.21 | **0.92 ± 0.12** | **n.d.** |
| **GFPgamma Ref(44)** | **3.33 ± 0.08** | **493** | **509** | 1.71 ± 0.35 | **0.78 ± 0.25** | **n.d.** |
| **Ivy Ref(43)** | **3.44 ± 0.06** | **500** | **515** | 0.98 ± 0.25 | **0.84 ± 0.12** | **n.d.** |
| **NowGFP (10s) Ref(45)** | **3.90 ± 0.09** | **492** | **502** | 0.79 ± 0.14 | **0.33 ± 0.13** | **n.d.** |

\*in contrast to the other maturation times that integrate protein folding and fluorophore maturation together, this value here only relates to fluorophore oxidation.

To further characterize the selected GFP variants we created yeast strains that endogenously expressed these proteins as C-terminal fusion to the cytoplasmic and nuclear localized yeast protein Ypd1. We then used these strains to compare different fluorescent proteins. In order to determine the photostability, we used a constant excitation intensity for all strains and acquired fluorescence intensity time traces for 45 s each. This revealed that NowGFP, the protein with the longest lifetime, was highly sensitive to photobleaching, whereas no major differences were detected between the other GFP variants (Fig. 1b). Western blotting revealed that they were expressed to similar levels, indicating that none of the proteins affected the expression of the fusion protein in a major way (Fig. 1c). For brightness comparison we quantified the concentration of different Ypd1-GFP fusions using FCS and used these measurements (Fig. 1d) to normalize the measured fluorescence intensities. This revealed that GFPgamma is by far the brightest GFP variant, while all the others exhibited similar molecular brightness (Fig. 1e,Table 1). It has to be noted that these values are only valid for the used system (excitation wavelength: 485 nm, major dichroic: zt 488/561 rpc, emission filter: ET525/50m), since the different GFP variants exhibit different excitation and emission optima (Table 1).

### Limits of sc-FLCCS examined by Monte-Carlo simulations

Molecular brightness, bleaching sensitivity and the fluorescence lifetime histograms are the basic characteristics of fluorescence proteins that contribute to the detection sensitivity of protein-protein interactions in sc-FLCCS. To explore how much each factor contributes to the overall performance of an individual fluorescent protein we used Monte-Carlo simulations and conducted virtual sc-FLCCS experiments. Thereby we simulated raw fluorescence photon events and generated simulated sc-FLCCS data. In contrast to real data, in simulated data the origin of individual photons is known. The simulated data was then correlated using conventional FCS analysis incorporating knowledge about the origin of the photons, or the data was correlated using sc-FLCCS approach, omitting the information about the origin of photons. The standard deviation of the first 15 points of the autocorrelation curve was used for comparison between FCS and sc-FLCCS. We first used this strategy to test the impact of molecular brightness. As expected, increased molecular brightness of fluorescent species improves the quality of the correlation curves in both approaches whereas lifetime filtering decreases the quality of the calculated correlation curves as indicated by an overall increased standard deviation for sc-FLCCS-derived curves (Fig. 2a). Next, we tested how the overlap of fluorescence lifetime histograms between the two different green fluorescent proteins affects the quality of the correlation curves (Fig. 2b). This revealed that the overlap has a significant impact, with a higher overlap worsening the quality of the correlation curves. To find out whether the impact of the lifetime filtering on the quality of the correlation curves does depend on the shape of the histograms we tested two different series; one series based on exponential histograms (inspired by fluorescence lifetime filtering) and a second series based on Gaussian distributed histograms (inspired by fluorescence spectral filtering(46)). This revealed no dependence on the shape of the histograms used for filtering (Fig. S3) and a sole dependence on the overlap of the area-normalized histograms.

**Fig. 2.**
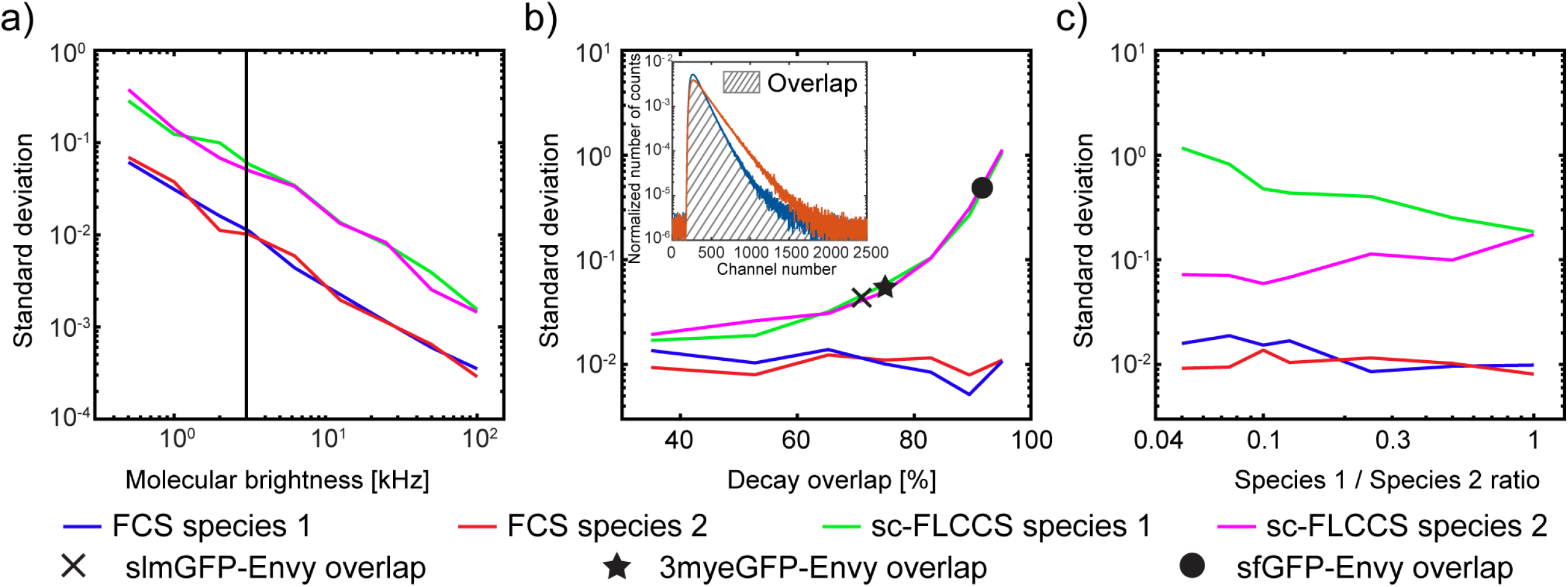
Quantification of the quality of resulting sc-FLCCS data using Monte-Carlo simulations. **(a)** Simulation of the impact of molecular brightness on the quality of autocorrelation curves - fluorescence lifetime histograms with 75% overlap. **(b)** The quality of the autocorrelation curves depends on the extent of overlap of the fluorescence lifetime histograms. Insert: Schematic illustration of the overlap of area normalized fluorescence lifetime histograms. **(c)** Higher difference in the species abundance decreases the data quality for the low abundant specie. Data in **(b)** and **(c)** are plotted for the molecular brightness of 3 kHz (indicated as a vertical line in **(a)**), which is a representative brightness of fluorescent proteins with our experimental conditions.

The quality of the correlation curves could depend on the relative abundance of the two fluorescent species. To address this, we varied the concentration of one species while keeping the concentration of the other constant (Fig. 2c). In the case of standard FCS and assuming no spectral cross talk at all, the concentration does not affect the results. In contrast sc-FLCCS showed a strong dependence on the relative abundance of species, with the best result for abundances of both species in the same concentration range. Together the analysis revealed that sc-FLCCS appears to be a valid alternative to dc-FCCS, but also that it has other intrinsic limitations imposed by the measurement principle.

### Testing of selected fluorescent proteins (FPs) for sc-FLCCS analysis

Our simulations demonstrate that the ability to separate correlation curves from different fluorophores with the same emission spectra depends on the difference in fluorescence lifetime histogram profiles, their molecular brightness and to some extent also their relative concentrations. To establish sc-FLCCS *in vivo*, we chose the two GFP variants with the shortest lifetime (3myeGFP and slmGFP) and tested each of them in combination with the three GFP variants with the longest lifetimes (Envy, GFPgamma and Ivy). We constructed strains where we expressed pairs of these GFP variants as N- and C-terminal fusion to the yeast protein Don1, which served as a spacer to keep the two fluorescent proteins with different lifetimes apart from each other(22). Next, we performed a sc-FLCCS measurement. For calculation of the fluorescence lifetime filters (see Materials & Methods) we used the strains that expressed individual Ypd1-FP fusion proteins (Fig. 1). This experiment revealed that the selected FP combinations provide data that is suitable for sc-FLCCS analysis (Fig. 3a). As expected from our simulations (Fig. 2b), the quality of the auto- and cross-correlation curves differs in terms of noise. Since both FPs are expressed as a part of the same translational unit, their concentrations and thus amplitudes of the auto-correlation functions should be the same. This is not the case in any of tested FP combinations presumably because of the differences in fluorophore maturation time and photo-stability (Fig. 3a, Table 1).

**Fig. 3.**
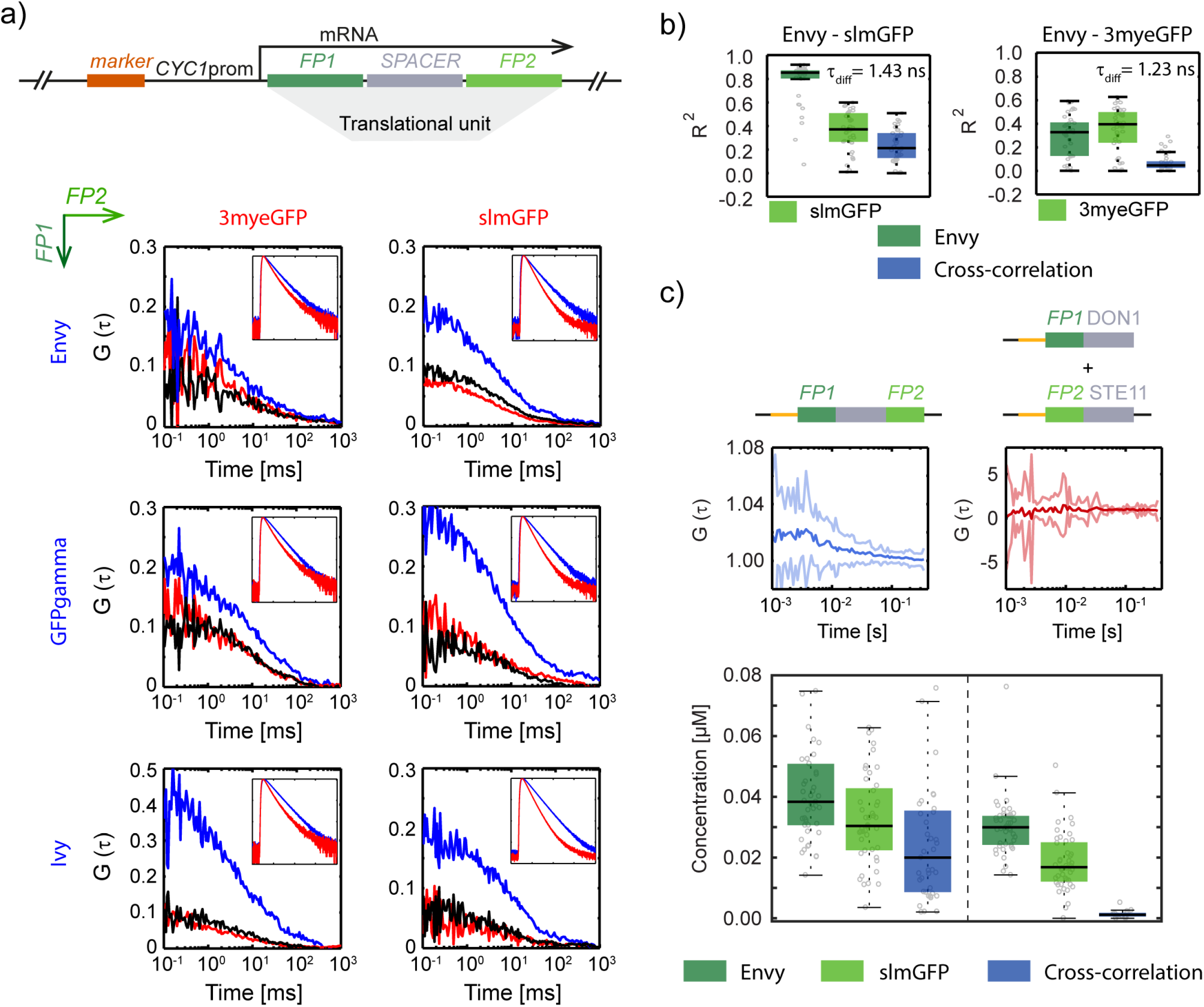
Proof of principle. **(a) Top:** Schematic illustration of the tandem fluorescent protein fusion, with N- and C-terminally tagged spacer protein (Don1). **Bottom:** Filtered auto- and cross-correlation functions were calculated for selected combinations of FPs. Fluorescence proteins in rows (blue lines) were attached N-terminally and proteins in columns C-terminally (red lines). Black lines correspond to the cross-correlation curves. Insets show corresponding fluorescence lifetime histograms (x-axis is number of channels ranging from 0-2000 and y-axis equate to normalized number of events ranging from 10^-4^ to 10^0^). **(b)** Comparison of the sc-FLCCS analysis for selected pairs of FPs as determined by R^2^ values. The higher the R^2^ value, the better the sc-FLCCS filtering and consecutive analysis. The difference in average fluorescence lifetimes between FPs (τ_diff_) is listed in the upper right corner of the corresponding box plot. **(c)** Comparison of the positive (left) and the negative (right) controls. **Top:** Schematic illustrations of the strains. Orange lines denote weak *CYC1* promoter. **Middle:** Cross-correlation curves, where solid dark line corresponds to the mean curve, light borders denote standard deviations. **Bottom:** Overall concentrations resulted from auto-correlation curves for each diffusing species and their complexes as determined by the sc-FLCCS analysis. Dashed line separates individual strains.

We further selected three combinations of FP pairs and tested them for sc-FLCCS application (Fig. 3a, Fig. S4). The selected pairs were: Envy/slmGFP (lifetime difference 1.43 ns, histograms’ overlap 71%), Envy/3myeGFP (lifetime difference 1.23 ns, histograms’ overlap 75%) and Envy/sfGFP (lifetime difference 0.73 ns, histograms’ overlap 92%). The quality of the resulted auto- and cross-correlation curves was quantified using the R^2^ value of the fit (Fig. 3b, Fig. S5 for Envy/sfGFP couple). The results correlate with the simulations, as shown by the black cross, black closed star and black closed circle in Fig. 2. This confirms the result from the simulations that the difference in the fluorescence lifetime histograms of the two used fluorescent proteins is a major factor influencing sc-FLCCS measurements. Because of the best resolution, the pair of slmGFP and Envy was chosen for further studies. In addition to the tandem fusion of both proteins (Envy-Don1-slmGFP) we also designed a negative control where both proteins are expressed independently (slmGFP-Don1 and Envy-Ste11) and no interaction was expected(22). We observed positive cross-correlation in the tandem fusion construct and no cross-correlation in the strain with independent expression units (Fig. 3c). This validates the concept of sc-FLCCS.

In summary, the best FP candidates for lifetime filtering are slmGFP, as the partner with the short fluorescence lifetime, and Envy, as the long fluorescence lifetime partner, both with reasonable brightness and good photo-stability.

### Challenging the limits of the sc-FLCCS analysis

Fluorescence resonance energy transfer (FRET) between two fluorophores depends on the spectral overlap, distance between the fluorophores and their mutual orientation. Because the FPs in sc-FLCCS are spectrally highly similar and might be spatially relatively close to each other we have to take into account the energy transfer. To test the impact of FRET, we prepared a construct where FPs are in a tandem in one translational unit (Fig. S6a). This is an extreme case where all the FPs are in the closest possible proximity, which is usually not the case for interacting tagged proteins. The fluorescence histograms were affected by the energy transfer and correct lifetime filtering was not possible. Thus, we obtained zero cross-correlation (Fig. S6b) and calculated concentrations were either overestimated (Envy) or underestimated (slmGFP, Fig. S6c; compare the values with Fig. 3c). This indicates that sc-FLCCS is sensitive to FRET. Using appropriate controls, e.g. specimen where only one of the components is tagged is required to ensure that there is no FRET occurring.

### Sc-FLCCS of proteasomal subunits

Next, we used sc-FLCCS to monitor the interaction between two proteins that are spatially well spaced so that FRET is unlikely to occur. We chose components of the proteasome, a large multi-subunit protease that functions as the major proteolytic activity in the cytoplasm of the cell, with functions in protein degradation and regulation. The proteasome consists of different substructures, i.e. the core particle that contains the active sites of the protease, and the base and the lid complexes that can dynamically interact with the core particle and that regulate access of ubiquitylated substrate proteins to the core particle (Fig. 4a)(47). We constructed strains where the core protein Pre6 was tagged with slmGFP and the lid protein Rpn7 was tagged with either Envy or 3mCherry. The distance between these proteins is about 12 nm, well beyond the range of FRET. We performed fluorescence fluctuation measurement and compared the results from sc-FLCCS (single color) and dc-FLCCS (dual color) analysis (Fig. 4b,c dashed lines separate different strains). First, we determined the overall concentrations of individual tagged proteins and quantified the amount of those which form the proteasome (Fig. 4b). We conclude that sc-FLCCS analysis of proteasomal proteins provide the same results as dc-FLCCS experiment. Considering R^2^ as a proxy for the quality of the data we found that the sc-FLCCS data could be less well fitted compared to the dc-FLLCS data. This indicates that filtering of photons in sc-FLCCS, which contains statistical uncertainty of photon origin, leads to data that is noisier than the dc-FLCCS data, where origin of each photon is known. Nevertheless, the sc-FLCCS yielded valid data demonstrating the applicability of the method. (Fig. 4c).

**Fig. 4.**
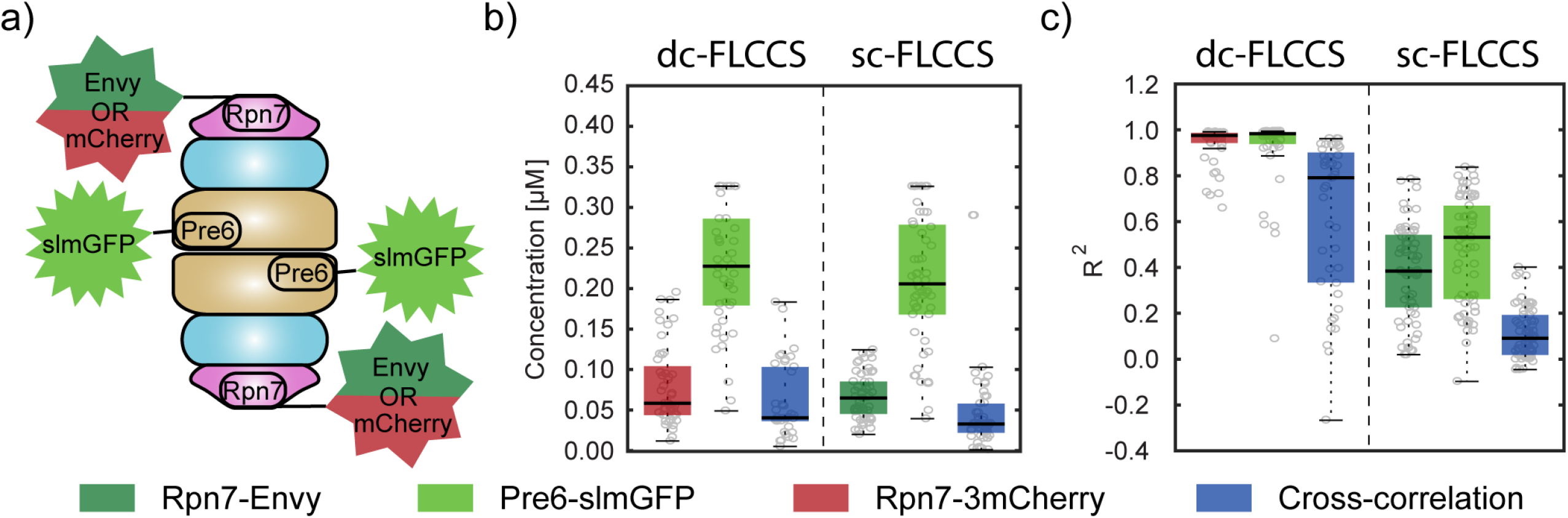
Interplay between Pre6 and Rpn7 examined by sc-FLCCS analysis. **(a)** Schematic organization of the yeast proteasome. Tagged proteins are indicated. Concentrations **(b)** and quality of the fit expressed by R^2^ values **(c)**. Overall concentrations resulted from auto-correlation curves for each diffusing species and their complexes as determined by the sc-FLCCS analysis. Dashed lines separate individual single- or multiple-tagged strains.

### Simultaneous monitoring of three proteasomal proteins using two channels

Measuring protein-protein interactions between three proteins inside the same cell is difficult. We decided to explore whether sc-FLCCS can be combined with dc-FLCCS. We chose the protein Rad23 as a third protein partner. Rad23 is a protein with an N-terminal ubiquitin like domain with several nuclear and cytoplasmic functions related to DNA damage repair and targeting of substrates to the proteasome(48–50). It has been reported that the E4 ubiquitin ligase Ufd2 and Rpn1, a component of the base part of the proteasome, compete for binding to Rad23. We used Rad23 tagged with 3mCherry, Pre6 with slmGFP and Rpn7 with Envy to performe a dual color sc/dc-FLCCS experiment. For comparison we used standard dc-FLCCS experiment, where only the interaction between Rad23 and Pre6 was monitored (Fig. 5a). The concentrations of individual proteins were converted to the scheme and the strength of the interaction was quantified by the dissociation constant (Fig. 5b). These results confirmed that dual color sc/dc-FLCCS enabled the detection of protein interactions between three different proteins of the proteasome using only one measurement in one strain.

**Fig. 5.**
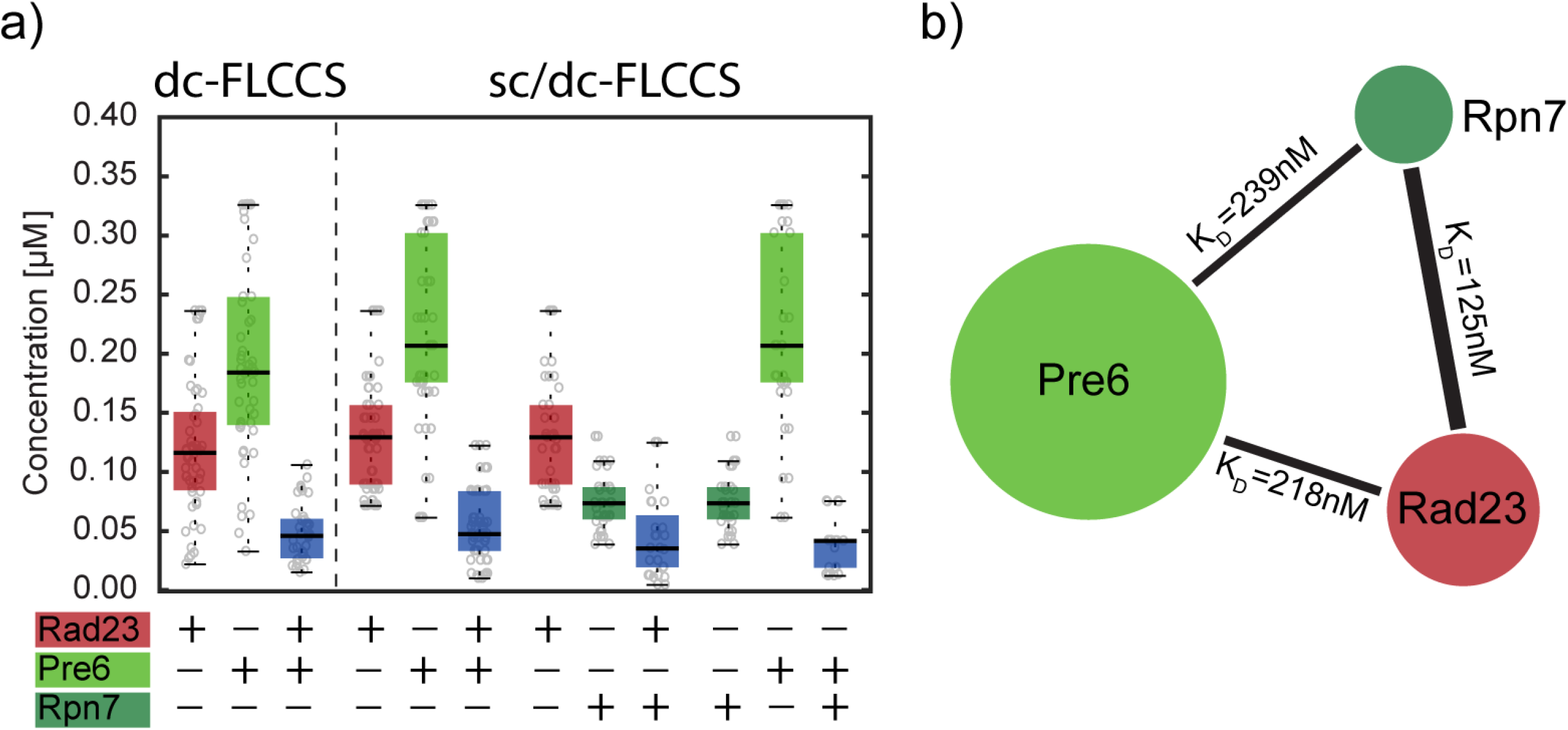
FLCCS analysis of a strain with three tagged proteasomal proteins. **(a)** Overall concentrations of individual proteins and concentrations resulted from cross-correlation. Dashed line separates different strains. The analysis of triple tagged strain required the combination of sc-FLCCS and dc-FLCCS (sc/dc-FLCCS). **(b)** Interaction map between Pre6, Rpn7 and Rad23. The size of the bubble illustrates the protein abundance and the thickness of the line between bubbles refers to the strength of the interaction between corresponding proteins.

## DISCUSSION

So far, sc-FLCCS(25, 31) was almost exclusively used for *in vitro* biophysical studies on model membrane systems(26, 27, 31, 51). There are also reports on sc-FLCCS studies performed in *in vivo*(52–54), but up to date it was not possible to perform fluorescence lifetime filtering on two different fluorescent proteins with matching spectral properties *in vivo*, either because of high similarity in their fluorescence lifetimes or because of insufficient photo-physical properties of those proteins with respect to photo-stability and molecular brightness.

To use fluorescent lifetime filtering as a mean to discriminate different fluorophores with the same spectral properties, we needed to identify GFP variants with optimal *in vivo* performance and extreme lifetimes, either very short or very long. From our experience, it is often not possible to derive these from published data, since these values were often determined *in vitro* or using different organisms. This is exemplified for NowGFP, which was reported as highly stable variant *in vitro* and *in vivo* in *E. coli, Drosophila* and mammalian cells(45, 55), but turned out to be highly photo-unstable in yeast, at least when imaged using our setup. This thus prompted us to evaluate several other green fluorescent proteins for sc-FLCCS applications ourselves.

We used Monte-Carlo simulations to explore the impact of different properties of fluorescence proteins on sc-FLCCS. We found that the critical parameter is indeed the difference in the fluorescence lifetime histograms of the used fluorescent proteins. A large overlap of the lifetime histogram means - in simple terms - that the uncertainty in photon assignment is too high for successful filtering To compensate for this loss, optimal performance parameters for other properties of fluorescent proteins are needed, in order to maximize the available overall ‘photon budget’, i.e. molecular brightness and photobleaching resistance. For further improvement, new fluorescent proteins with longer lifetimes would be needed. Alternatively, applications where one of the GFPs is replaced by chemical dye labeling, e.g. using SNAP or HALO tags(56, 57), should be possible in organisms where such dye labeling strategies are feasible (which is not easily the case in *S. cerevisiae*). Chemical dyes exhibit a much broader range of physico-chemical properties, including much longer lifetimes(58). Thereby, much more precise “photon assignment” can be reached, thus enabling ‘perfect’ sc-FLCCS applications.

Our results demonstrate that sc-FLCCS analysis is an alternative and valid approach to the standard dual color FCCS or dc-FLCCS methods. The advantage of the sc-FLCCS over standard dual color methods is the reduction of two excitation wavelengths to only one while being able to resolve two spectrally similar fluorescence proteins. This is based on their differences in fluorescence lifetime histograms, but it also requires that these are determined for individual fluorescence proteins prior the sc-FLCCS analysis. Moreover, the fluorescence lifetime does not depend on the protein abundance, excitation wavelength, filter configuration and photo-bleaching, which is a definite advantage of this method.

A disadvantage of sc-FLCCS however, apart from the ‘increased uncertainty due to filtering’ is the sensitivity of the method to FRET, which affects the lifetime histograms, impedes the analysis, and essentially makes the method useless in the case of two tightly interacting small proteins (Fig. S6c). However, the FRET efficiency decays with the 6^th^ root of the distance which makes sc-FLCCS suitable especially for large complexes. It is important to note that FRET does also affect dual color FCCS. In the case of EGFP and mCherry, typical green and red FPs used in dual color FCCS, which have highly overlapping emission (EGFP) and excitation spectra (mCherry), it causes an artificial underestimation (green channel) and overestimation (red channel) of the concentration of the individual fluorophores and it also affects the amount of interaction derived from cross-correlation. This fact is usually not considered when performing classical dual color FCCS experiment *in vivo*. An additional disadvantage of sc-FLCCS over standard dc-FLCCS is the requirement that both concentrations shall be in the same range, at least for situations with two fluorescent proteins with a significant lifetime histogram overlap.

Nevertheless, in our proof of concept experiment we have demonstrated that sc-FLCCS in combination with dc-FLCCS allows to distinguish and quantify the interaction of three proteins, measured simultaneously in one strain in one measurement using an instrument set up for two wavelengths only. Further improvements of sc-FLCCS towards four interaction partners is also thinkable, given the existence of new variations of red fluorescence proteins with prolonged fluorescence lifetime (e.g. mScarlet and its variants(59)). Combination of mScarlet (fluorescence lifetime 3.9 ns) and mCherry (fluorescence lifetime 1.5 ns) would be a perfect RFP-like couple for sc-FLCCS experiment in the red spectra. Thus, dual color sc-FLCCS (sc in green + sc in red channels) could provide the information about four different proteins in just one measurement. In classical 2-color dc-FLCCS experiment one would need pair wise measurements which would require 6 strains to measure all interactions. Using dual color sc-FLCCS decreases significantly the necessary measurement time when determining the interaction map of multiple proteins *in vivo*.

In summary, we have presented a possibility to use single color FLCCS to analyze the interaction of two proteins *in vivo* and we have explored and discussed the advantages and disadvantages when compared to dc-FLCCS. Our *in vivo* measurements of important ‘vital’ parameters of many green fluorescent protein variants furthermore enables the selection of best couple of GFP-like fluorescence proteins not only for sc-FLCCS, but also for potential applications such as fluorescence lifetime imaging, and we demonstrated the usefulness of the sc-FLCCS method in determination of the concentrations and intermolecular interactions in yeast cells.

## AUTHOR CONTRIBUTIONS

MS and MK designed experiments. MS, KH and MR performed experiments and processed data. AB performed Monte-Carlo simulations. MS and MK wrote the paper.

## ACKNOWLEDGMENT

The authors thank to Dr. Alexander S. Mishin for providing us with the NowGFP containing plasmids. We acknowledge support from the German Research Foundation (DFG grant INST 35/1133-1 FUGG). MS was supported by fellowships from Cellnetworks and the Alexander von Humboldt Foundation. MS also thanks to the Czech Science Foundation (grant 19-08304Y). AB acknowledges ERD Fund-Project No. CZ.02.1.01/0.0/0.0/16_013/0001775.

